# Cortical reactivations predict future sensory responses

**DOI:** 10.1101/2022.11.14.516421

**Authors:** Nghia D. Nguyen, Andrew Lutas, Jesseba Fernando, Josselyn Vergara, Justin McMahon, Jordane Dimidschstein, Mark L. Andermann

**Affiliations:** Program in Neuroscience, Harvard University, Boston, MA 02215, USA; Division of Endocrinology, Department of Medicine, Beth Israel Deaconess Medical Center, Boston, MA 02215, USA; Stanley Center for Psychiatric Research, Broad Institute of Harvard and MIT, Cambridge, MA 02142, USA; Diabetes, Endocrinology, and Obesity Branch, National Institutes of Diabetes and Digestive and Kidney Diseases, National Institutes of Health, Bethesda, MD 20892, USA

**Author notes:** **Materials and correspondence** Correspondence and requests for materials should be addressed to Mark L. Andermann.

## Abstract

Prevailing theories of offline memory consolidation posit that the pattern of neurons activated during a salient sensory experience will be faithfully reactivated, thereby stabilizing the entire pattern^1-3^. However, sensory-evoked patterns are not stable, but instead drift across repeated experiences^4-7^. To investigate potential roles of reactivations in the stabilization and/or drift of sensory representations, we imaged calcium activity of thousands of excitatory neurons in mouse lateral visual cortex. Presentation of a stimulus resulted in transient, stimulus-specific reactivations during the following minute. These reactivations depended on local circuit activity, as they were abolished by local silencing during the preceding stimulus. Contrary to prevailing theories, reactivations systemically differed from previous patterns evoked by the stimulus. Instead, they were more similar to future patterns evoked by the stimulus, thereby *predicting* representational drift. In particular, neurons that participated more or less in early reactivations than in stimulus response patterns subsequently increased or decreased their future stimulus responses, respectively. The rate and content of these reactivations was sufficient to accurately predict future changes in stimulus responses and, surprisingly, the decreasing similarity of responses to distinct stimuli. Thus, activity patterns during sensory cortical reactivations may guide the drift in sensory responses to improve sensory discrimination^8^.

## Introduction

In the absence of salient ongoing sensory stimuli, the brain may instead learn from prior experiences by repeatedly replaying or reactivating neural patterns that were active during past experiences^1, 9-14^. Such reactivations involve temporally condensed, hyper-synchronous events that occur during quiet waking and sleep^1, 11-13^. First observed and most commonly studied in the hippocampus, reactivations have also been observed in the amygdala, prefrontal cortex, visual cortex, striatum and elsewhere^15-29^.

Reactivations, by definition, are patterns of activity that are similar to those that occurred during recent experiences^1, 30, 31^. However, in part due to the limited recording of tens to hundreds of neurons in prior studies, it remains unclear whether reactivations are faithful copies of activity patterns that occurred during prior experiences. Alternatively, given that stimulus response patterns gradually change across trials (i.e. representational drift^4-7, 32^), reactivations might instead more closely resemble future responses to the same stimulus. To more accurately compare the content and dynamics of stimulus response patterns and offline reactivations across trials, we recorded the activity of ∼6,600 neurons simultaneously in lateral visual cortex while mice passively viewed well-controlled presentations of the identical stimuli, separated by long inter-stimulus intervals where reactivations should occur^29, 33^.

## Results

### Distributed reactivations of stimulus-evoked response patterns in lateral visual cortex

Awake, head-fixed mice (n = 4) passively viewed 64 presentations per day of each of two visual stimuli across daily sessions during cellular imaging (Fig. 1a; Stimulus 1 [S1] and Stimulus 2 [S2], presented in random order; 2-s duration; 11 ± 1 sessions per mouse, mean ± SEM). Unlike conventional sensory mapping protocols, each presentation was followed by a long 58-s inter-trial interval (ITI) to assess for possible offline reactivations of stimulus-evoked response patterns (Fig. 1a). To track neural activity in glutamatergic neurons throughout lateral visual cortex, we made multiple viral injections of a genetically-encoded calcium indicator (Cre-dependent expression of jGCaMP7s^34^ in Emx1-Cre^35^ mice; Fig. 1b). First, we defined visual cortical areas using a brief epifluorescence imaging protocol^36^. We then combined sensory stimulation with widefield two-photon Ca^2+^ imaging to simultaneously record activity of many thousands of distinct neurons (6,623 ± 166 neurons/session, mean ± SEM across 42 sessions from 4 mice) across three planes within layers II-IV of four lateral visual cortical areas (Fig. 1b). For each neuron, we defined the activity time course as the deconvolution of the neuropil-corrected Ca^2+^ fluorescence trace^37^ (normalized, for each neuron, to the mean of the top 1% of transient calcium events). We focused analyses on stimulus-driven neurons (1,517 ± 134 neurons/session, mean ± SEM), which either showed a preferential increase in activity to S1 or S2, or responded similarly to both (Fig. 1c).

**Figure 1.**
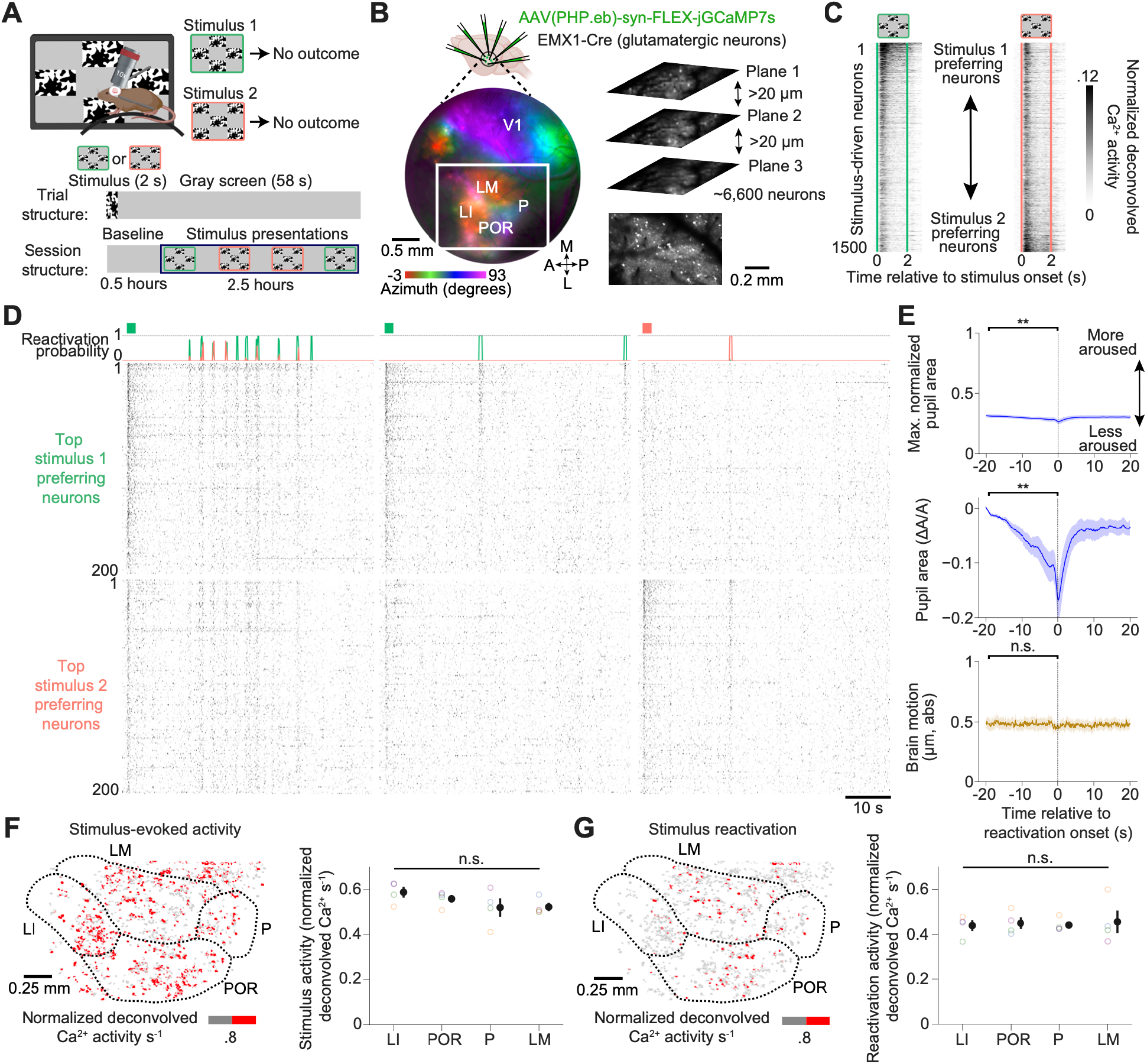
Distributed stimulus reactivations in lateral visual cortex during quiet waking. **a**, Setup for two-photon imaging of awake, head-fixed mice during repeated, passive presentation of two checkerboard patterns containing binarized white noise (Stimulus 1 [S1] or Stimulus 2 [S2]). Following a 0.5 hour baseline period, the stimuli (2-s duration, 58-s inter-trial interval) were presented in random order across 2.5 hours. **b**, Top: Cre-dependent jGCaMP7s expression in glutamatergic neurons via local injection at six sites across visual cortex in Emx1-Cre mice. Bottom: epifluorescence retinotopic mapping of local preference for specific locations in visual space identified several visual areas: V1: primary visual cortex; LM: lateromedial; POR: postrhinal; LI: laterointermediate; P: posterior. Simultaneous imaging of ∼6,600 neurons in layers II-IV of visual association cortex (white rectangle) across 3 vertical planes using two-photon Ca^2+^ imaging. **c**, Trial-averaged, deconvolved peri-stimulus Ca^2+^ activity from an example session. Some stimulus-driven neurons responded preferentially to S1 or S2 (top and bottom) while others responded equally to both (middle). **d**, Raster plot of ongoing deconvolved Ca^2+^ activity of the top 200 S1-driven and S2-driven neurons during and following three example stimulus presentations.

As illustrated in three example trials, we observed many events consisting of transient (<1 s) moments of synchronous activity of stimulus-driven neurons in the tens of seconds following stimulus presentation (Fig. 1d). Using a multinomial logistic regression-based classifier, we assigned a probability that these synchronous events resembled response patterns during S1, S2, or neither (Fig. 1d; see Extended Data Fig. S1a and Methods for details). Synchronous ongoing activity patterns that resembled S1 or S2 response patterns with high classifier probability (> 0.75) were over three times less frequent in data in which the identity of each neuron was shuffled, or in which the set of neurons that defined synchronous ITI events was shuffled to include neurons not driven by S1 or S2 (Extended Data Fig. S1b). Thus, we defined S1 or S2 stimulus reactivations as synchronous ITI events with classifier probabilities greater than 0.75 for either S1 or S2, respectively. We confirmed the similarity of these stimulus reactivations to stimulus-evoked response patterns by grouping neurons based on their mean response magnitude during stimulus presentations (Extended Data Fig. S1d). As expected, the most S1- and S2-driven neurons were selectively active during S1 and S2 reactivations, respectively, while more weakly stimulus-driven neurons and non-driven neurons were not selectively active.

We wondered whether imaging thousands of neurons instead of hundreds (cf. Sugden et al., 2020) increased our sensitivity for capturing stimulus reactivations. Indeed, when only using a random 10% of neurons instead of the full dataset of ∼6,600 neurons per session, over two-thirds of identified reactivations were missed, and the rate of false-positive reactivations also increased (Extended Data Fig. S1e-f). The neurons that contributed to stimulus reactivations were distributed across the four lateral visual cortical areas that we imaged simultaneously, with similar levels of activity in each cortical area during both stimulus presentations and stimulus reactivations (Fig. 1f-g).

Reactivations were associated with moments of particularly low arousal (Fig. 1e). Seconds before the onset of a stimulus reactivation, pupil area – an index of arousal^38^ – was already at ∼1/3 of that observed during active movement, and briefly constricted further during reactivations. Reactivations were not accompanied by increased body movement (approximated using brain motion) or eye movement (Fig. 1e and Extended Data Fig. S1c).

### Reactivations rates are biased by the previous stimulus and decline with experience

The content of awake reactivations of spatiotemporal patterns of activity (i.e. replay) in the hippocampus can be biased towards salient (novel, rewarding or aversive) recent experiences^14, 23, 24, 39-42^, with a reactivation rate that often decays across trials^22, 39^. As illustrated for an example session (Fig. 2a) and quantified below, cortical sensory reactivations exhibited similar properties.

**Figure 2.**
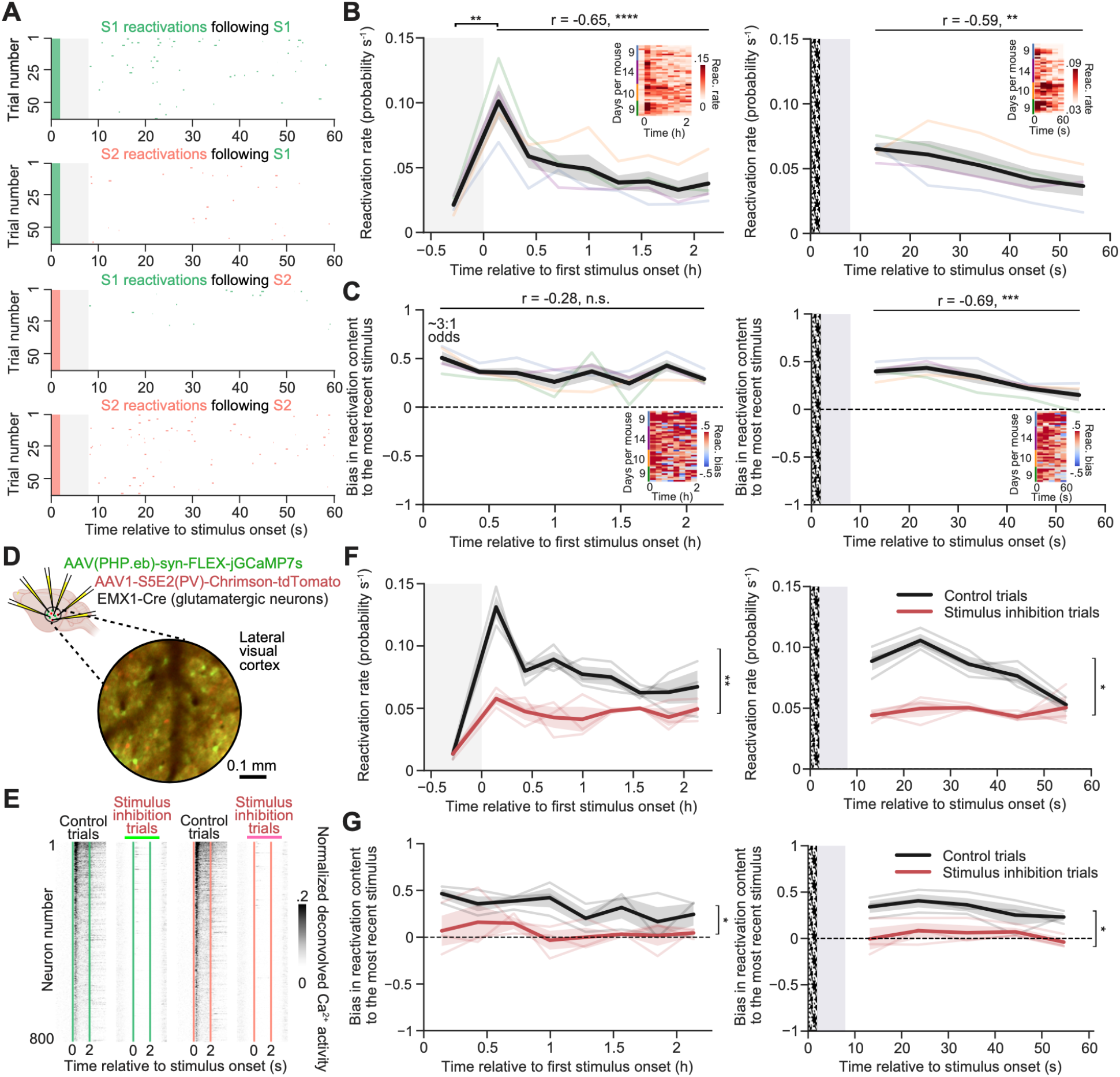
Cortical responses to stimuli drive subsequent reactivations. **a**, Example single-session raster plot of S1 and S2 reactivations (green and red dots) following the presentation of S1 (top) or S2 (bottom). **b**, Left: reactivation rates (sum of probabilities of S1 or S2 reactivations) across the session, including the 0.5-hour baseline period prior to any stimulus presentations (shaded region; n = 4, two-tailed paired t-test, two-sided linear least-squares regression). Colored lines denote mean per mouse across sessions. Black line denotes mean across all mice. Inset: heat map of reactivation rate for each daily session and mouse. Right: reactivation rate during the inter-trial interval (n = 4, two-sided linear least-squares regression). Inset: heat map of reactivation rates for each daily session and mouse. Shaded region: excluded portion of inter-trial interval. White-noise bar: stimulus presentation. **c**, Left: bias index of reactivation content (positive values indicate bias towards the most recent stimulus, see Methods; n = 4, two-sided linear least-squares regression). Inset: heat map illustrating bias for each daily session and mouse. Right: bias throughout the inter-trial interval (n = 4, two-sided linear least-squares regression). Inset: heat map of bias for each daily session and mouse. **d**, Schematic and mean *in vivo* image of selective viral expression throughout visual cortex (6 injection sites) of jGCaMP7s and Chrimson in glutamatergic neurons (Emx1-Cre) and parvalbumin interneurons (due to S5E2 enhancer), respectively. **e**, Mean stimulus-evoked activity of driven neurons across trials from one example session. On a random set of ‘stimulus inhibition trials’ involving S1 or S2 presentation (vertical green and red lines), stimulus-evoked activity was suppressed from 1-s prior to 1-s after stimulus presentation (light green and pink bars) by pulsing 10 mW red light for 4 ms at 16 Hz. **f**, Left: reactivation rates across the session, averaged across control vs. stimulus inhibition trials (n = 3 mice). Right: reactivation rates throughout the inter-trial interval (n = 3 mice). **g**, Left: reactivation bias on control vs. stimulus inhibition trials (n = 3 mice). Right: reactivation bias throughout the inter-trial interval (n = 3 mice). **f-g**: light gray/red lines: mean per mouse across sessions. Black and red lines: mean across mice. Comparisons: two-tailed paired t-tests between means of the two traces.

We first evaluated changes in the rate of stimulus reactivations throughout a session across mice. Reactivation rates increased ∼4-fold above baseline levels following the first stimulus presentation and decayed back to baseline over two hours (Fig. 2b; post-stimulus reactivations were also longer in duration than during the baseline period, Extended Data Fig. S2a)^43^. Within the inter-trial interval following each stimulus presentation, stimulus reactivation rates increased and then decayed across tens of seconds (Fig. 2b). The *content* of reactivation events following a stimulus showed a strong, 3:1 bias towards those resembling the most recent stimulus – an effect that persisted throughout each session but that became weaker over tens of seconds following each stimulus presentation (Fig. 2c).

Reactivation rates appeared to scale inversely with the frequency of recent exposures to a stimulus, suggesting an effect of stimulus salience. This effect was not only evident in the gradual decline in reactivation rates across the session (Fig. 2b), but also in the increased rate of reactivations when the previous stimulus was preceded by a different stimulus versus by the same stimulus (Extended Data Fig. S2b).

These rate and bias effects were not due to changes in overall levels of ongoing activity, which remained stable across hours and throughout each ITI (Extended Data Fig. S2c). Furthermore, these effects could not be explained by a change in global arousal state (indexed by pupil size). Arousal was stable across hours and throughout each ITI (Extended Data Fig. S2d). Of note, the strong rate and bias effects described above were far less evident when only analyzing hundreds rather than thousands of simultaneously recorded neurons (Extended Data Fig. S2e).

### Local sensory-evoked activity is required for subsequent reactivations

We wondered whether the rate and bias effects described above required the same cortical neurons that participate in reactivations to be active during the preceding stimulus presentation. To test this, we optogenetically silenced stimulus-evoked activity in excitatory neurons throughout the imaged region of lateral visual cortex. To this end, we made local viral injections of the red-shifted opsin Chrimson under the S5E2 enhancer, which selectively targets parvalbumin interneurons^44^, along with Cre-dependent jGCaMP7s in Emx1-Cre mice (Fig. 2d and Extended Data Fig. S2f). We directly compared reactivation rates on control trials (used to train the classifier) vs. trials with photostimulation of parvalbumin interneurons (50% of trials selected at random). Photostimulation inhibited >90% of peri-stimulus activity of excitatory neurons in lateral visual cortex (Fig. 2e and Extended Data Fig. S2g), but did not affect arousal (Extended Data Fig. S2h). Further, it did not affect cortical activity levels during the ITI, which gradually returned to baseline within seconds following photosilencing (Extended Data Fig. S2i).

Inhibition during stimulus presentation strongly reduced subsequent stimulus reactivation rates (Fig. 2f, n = 3 mice, 8 ± 1 sessions per mouse, mean ± SEM). Strikingly, this inhibition also abolished the subsequent bias in the content of stimulus reactivations towards the stimulus presented (Fig. 2g, n = 3 mice). Therefore, local activity during a sensory experience is necessary for the subsequent appearance of biased reactivations in lateral visual cortex.

### Lateral visual cortex orthogonalizes stimulus representations

Having characterized visual cortical reactivations, we next considered potential functional roles for these events. Previous studies of hippocampus suggests that reactivations of certain experiences may play a role in neural plasticity and learning^1-3^. While visual cortical response patterns are known to gradually change across repeated presentations (i.e. representational drift^4-7, 32^), how reactivations might shape this plasticity process remains unknown.

To investigate these changes in stimulus response patterns, we first considered single-trial stimulus-evoked responses averaged across all neurons. This response was constant across repeated stimulus presentations, consistent with some previous studies^45-48^ (Fig. 3a). We next considered whether evoked response *patterns* might change across repeated presentations. Inspection of single-trial responses from a typical example session suggested that many neurons responded initially to both stimuli, but gradually lost their responses to one or the other stimulus (Fig. 3b). As such, the patterns of population responses to the two stimuli should become more dissimilar across repeated presentations, potentially facilitating stimulus discriminability. We quantified this phenomenon using the running Pearson’s correlation between neighboring S1 and S2 single-trial response patterns. This correlation decreased throughout the session, indicating that the two stimulus representations became more orthogonal (i.e. decorrelated) across trials (Fig. 3c).

**Figure 3.**
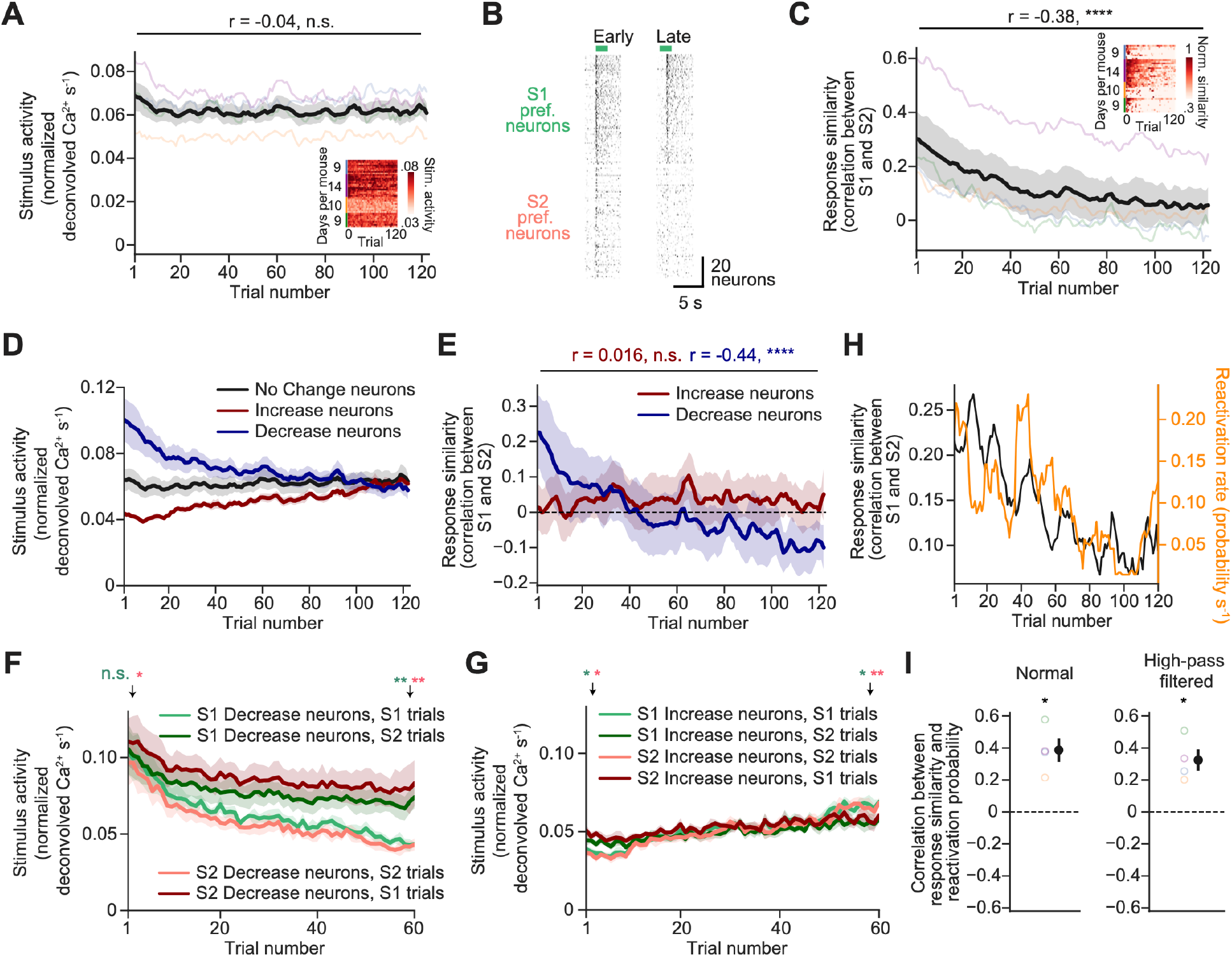
Progressive separation of stimulus response patterns correlates with reactivation rates. **a**, Mean stimulus-evoked activity per trial (across all S1 and S2 trials), averaged across all stimulus-driven neurons. Traces were smoothed with a 3-trial sliding average. Colored lines indicate mean across days per mouse. Black line indicates mean across mice (n = 4, two-tailed least squares linear regression). Inset: mean activity for each day and mouse. **b**, Raster plots of activity of S1- and S2-preferring neurons during two S1 trials, early and late in a session. **c**, Response similarity (running correlation between responses of neighboring S1 and S2 trials). Traces were smoothed with a 3-trial sliding average. Colored lines indicate mean across days per mouse. Black line indicates mean across mice (n = 4, two-tailed least squares linear regression). Inset: heat map of the peak-normalized response similarity across all days and mice. **d**, Same as mean trace in **a** but calculated separately for neurons whose stimulus responses increase, decrease, or show no change across trials (n = 4). **e**, Same as mean trace in **c** but calculated separately using only neurons whose stimulus responses increase or decrease across trials (n = 4, two-tailed least squares linear regression, Holm-Bonferroni corrected). **f**, Evoked activity of Decrease neurons in **d**, plotted separately for neurons whose responses decrease to S1 (light green) or S2 (pink). Also shown are the responses to the other stimulus for these two sets of neurons (dark green and dark red, respectively; n = 4, two-tailed paired t-test within each set, Holm-Bonferroni corrected). **g**, Same as **f** but for Increase neurons. **h**, Example response similarity vs. reactivation rate across all trials for a single day. **i**, mean correlation (colored circles: per mouse, averaged across sessions; black disc: mean ± SEM across mice) between the two variables for unfiltered traces (left) and after high-pass filtering each trace to examine more rapid co-fluctuations (n = 4, two-tailed t-test vs. 0, Holm-Bonferroni corrected). Traces were smoothed with an 8-trial sliding average.

To determine which neurons contributed to stimulus orthogonalization, we considered groups of neurons based on their change in stimulus-evoked activity from early to late trials. We defined Increase neurons as those that exhibited an increase in evoked activity from early to late trials, while Decrease neurons showed the opposite trend, and No Change neurons did not change evoked activity across the session (see Methods, Fig. 3d, Extended Data Fig. 3a). When we quantified the changes in correlation between S1- and S2-evoked response patterns across trials separately for the subsets of Decrease or Increase neurons alone, we observed a similar orthogonalization of stimulus response patterns within the set of Decrease neurons but not within the set of Increase neurons (Fig. 3e; by definition, responses patterns across the set of No Change neurons remained unchanged, Extended Data Fig. S3b). The key role of Decrease neurons in driving overall population orthogonalization was confirmed by analyses showing less overall population orthogonalization when removing Decrease neurons vs. Increase neurons (Extended Data Fig. S3c). To visualize the activity dynamics of neurons whose responses to S1 or S2 decreased across trials (S1 Decrease or S2 Decrease neurons, respectively), we plotted stimulus-evoked activity across both S1 and S2 trials (Fig. 3f). S1 Decrease neurons decreased their responses to S2 less than to S1, thereby increasing their response tuning to S2 (Fig. 3f). Similar effects were observed in S2 Decrease neurons (Fig. 3f). Consistent with the weaker across-trial orthogonalization in the set of Increase neurons across presentations (Fig. 3e), these neurons showed more uniform increases in activity across trials during both S1 and S2 across trials (Fig. 3g). Thus, while the presence of both Increase neurons and Decrease neurons results in consistently balanced mean responses across all neurons, Decrease neurons are largely responsible for stimulus orthogonalization in lateral visual cortex.

Consistent with the sharp drop across the session in both the similarity in population responses to S1 vs. S2 (Fig. 3c) and in stimulus reactivation rates (Fig. 2b) across the session, these two measures were positively correlated across the session (Fig. 3h-i). Interestingly, even after removing slow drift in these two measures across the session, they co-fluctuated at faster time scales (∼10 trials; Fig. 3i; correlation peaks at zero-trial delay, Extended Data Fig. S3d). Therefore, the evolution of the representation of a stimulus in lateral visual cortex across trials tightly correlates with the rate of stimulus-specific reactivations of that stimulus, suggesting a possible role for reactivations in driving gradual changes in stimulus response patterns and their orthogonalization.

### Early stimulus reactivations predict late stimulus-evoked patterns

To better understand the relationship between reactivation patterns and the changes in single-trial stimulus-evoked response patterns across a session, we projected each pattern onto the axis of population activity between early and late stimulus-evoked responses within a session (i.e. dot product between the activity patterns and the vector difference between the mean stimulus-evoked response patterns on early trials and on late trials within a session; Fig. 4a,b). Both S1- and S2-evoked response patterns showed progressive changes from early to late trials (i.e. representational drift^4-7, 32^), with larger changes early in each session (Fig. 4b and Extended Data Fig. S4a). If stimulus reactivations were a copy of the previous stimulus-evoked response pattern, as suggested in previous studies^1, 11, 12^, we would expect a similar evolution of stimulus reactivation patterns from early to late trials. However, the projected stimulus reactivation patterns were instead stable across trials (Fig. 4b and Extended Data Fig. S4a). Strikingly, even after the first stimulus presentation, this projected pattern of stimulus reactivations already resembled the pattern of stimulus-evoked responses late in the session (Fig. 4b and Extended Data Fig. S4a). This finding was not due to the differential influence of early vs. late trials on classifier sensitivity, as we separately trained classifiers on early, middle, and late portions of a session, and used the highest of the three estimated matching probabilities at each time point (see Extended Data Fig. S1a and Methods).

**Figure 4.**
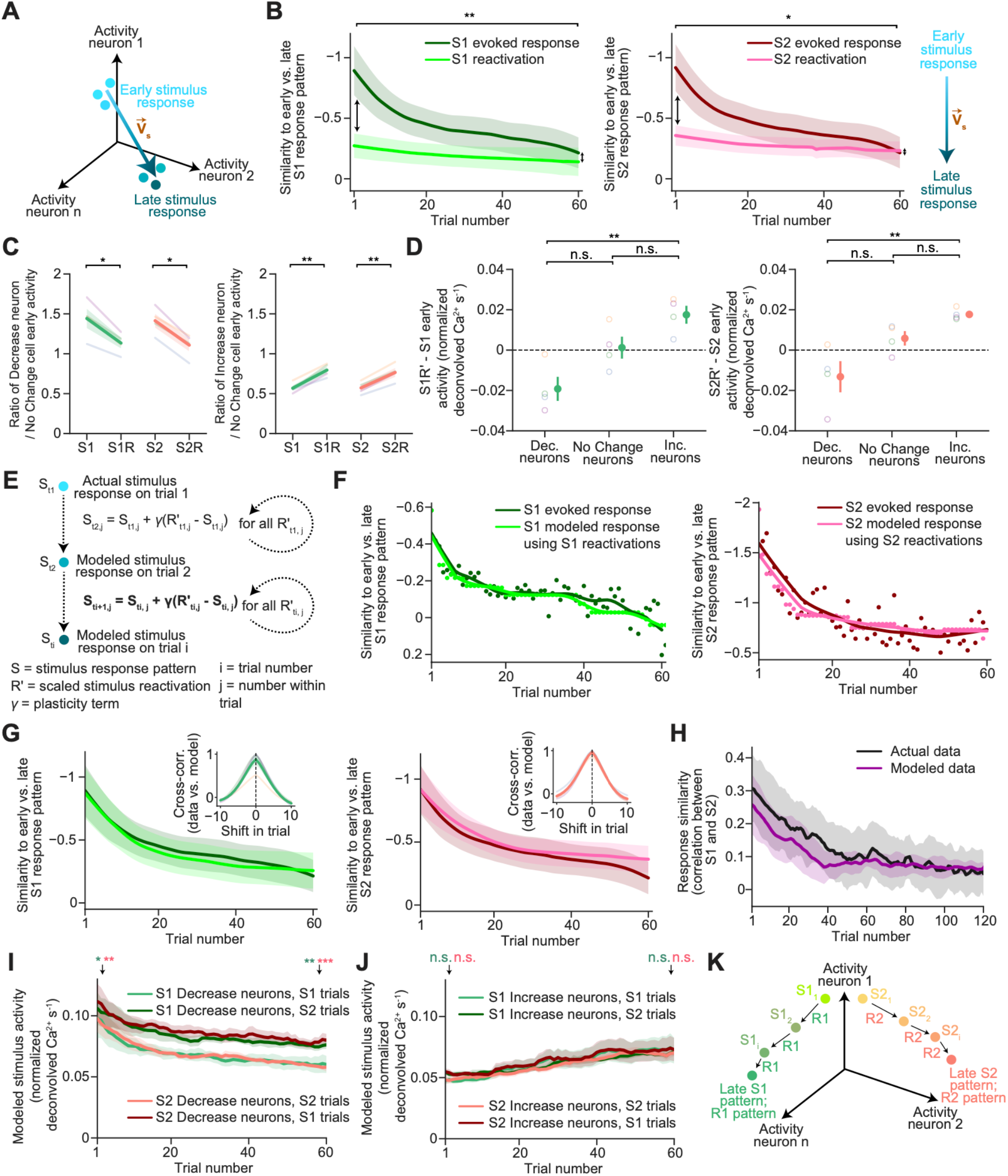
Reactivations predict future stimulus responses. **a**, Schematic of the evolution of single-trial stimulus response patterns across the session in high dimensional space, where each axis reflects the stimulus-evoked activity of one stimulus-driven neuron. V_s_ denotes the vector along the axis from early to late response patterns. **b**, For each trial, we projected the response pattern (during S1 or S2) and any stimulus reactivations during the inter-trial interval (S1R or S2R) onto the axis between early and late stimulus-evoked response patterns within a session (i.e. onto V_s_; mean across all days from 4 mice). **c**, Left: ratio of mean activity across trials early in each session in Decrease neurons relative to No Change neurons during S1 presentations vs. S1R events, and during S2 vs. S2R events (two-tailed paired t-test, Holm-Bonferroni corrected; colors indicate individual mice). Right: same but for Increase neurons. **d**, Difference between a neuron’s peri-stimulus activity and peri-reactivation activity early in each session, averaged across neurons in each class. Prior to neuron-by-neuron subtraction, reactivation patterns were scaled (by ∼1.4x; denoted R) to have similar mean activity as stimulus-evoked patterns. **e**, Model using reactivations to predict future stimulus responses. Beginning with the actual stimulus response pattern on the first trial, we iteratively modeled changes to subsequent stimulus responses by adding the difference between each scaled reactivation pattern during the intervening inter-trial interval and the previous stimulus response pattern, multiplied by a single free parameter, γ. **f**, Comparison of projection of the actual stimulus-evoked response patterns in **b** (dark green/red dots and smoothed trace) with the modeled patterns (light green/pink dots and smoothed trace), projected onto V_s_ for a single session. **g**, Same as **f** but mean across all days and mice (n = 4). Insets: cross-correlation between high-pass filtered actual and modeled projections of stimulus-evoked response patterns. **h**, Response similarity as measured by the correlation between the mean response patterns during nearby S1 and S2 trials, plotted for actual and modeled stimulus responses (n = 4 mice). Traces were smoothed with a 3-trial sliding average. **i**, As in **Fig. 3f**, mean stimulus-evoked activity using modeled data, averaged across S1 Decrease or S2 Decrease neurons (sets are defined using actual data; cf. **Fig. 3d,f**; n = 4, two-tailed paired t-test, Holm-Bonferroni corrected). **j**, same as **i** but for Increase neurons (cf. **Fig. 3d,g**). **k**, Conceptual model. S1- and S2-evoked response patterns are similar early in each session. With each stimulus-induced reactivation, S1 and S2 response patterns are pulled towards their respective reactivation pattern. This is sufficient to predict which neurons will increase or decrease their responses over trials, as well as the rate of change and orthogonalization of the stimulus-evoked responses.

To determine which neurons contributed to the difference between early stimulus-evoked patterns and early reactivation patterns, we compared the mean activity across Decrease or Increase neurons relative to the mean activity across No Change neurons. Even during early trials within a session, Decrease neurons showed relatively less activity during reactivations than during stimulus-evoked responses, while the opposite was true for Increase neurons (Fig. 4c). To compare the activity of individual neurons more directly, we scaled up activity levels across all neurons during early stimulus reactivations by a common scale factor (1.4x; see Methods) such that the mean activity across the population was roughly similar to that observed during early stimulus-evoked responses (Extended Data Fig. S4b). When we then subtracted early stimulus-evoked responses from these scaled early stimulus reactivations, we again found that Decrease neurons were relatively less active during early reactivations than during early stimuli, while the converse was true for Increase neurons (Fig. 4d). Further, No Change neurons showed similar levels of participation in early stimulus responses and reactivations (Fig. 4d). Together, these data show that neurons that participate in early reactivations below, at, or above their level of participation in early stimulus-evoked responses will subsequently decrease, show no change, or increase their stimulus-evoked activity later in the session, respectively. This suggests that both the rate and pattern of stimulus reactivations could be important in guiding the dynamics and nature of the change in stimulus-evoked response patterns, rather than simply stabilizing the previous stimulus-evoked response pattern.

### Stimulus reactivations model future stimulus representations

If stimulus reactivations did guide future changes in stimulus representations, then stimulus reactivations alone might be sufficient to predict the nature and rate of change in the patterns of future stimulus-evoked responses. We developed a simple heuristic model that uses only the stimulus-evoked response pattern on the first trial and all stimulus reactivations observed on every trial to iteratively predict stimulus-evoked response patterns on the second through the last trial (Fig. 4e). In this model, each time a S1 stimulus reactivation occurs following a S1 trial (and likewise for S2), we modified the estimate of the upcoming stimulus-evoked response pattern by adding the difference between the scaled reactivation and the current stimulus-evoked pattern, multiplied by a fixed plasticity term (Fig. 4e). We parametrically varied this plasticity term to find the best fit value and applied the same single value across all sessions and mice (Extended Data Fig. S4c). Intuitively, this model should drive faster changes in the stimulus-evoked response pattern early in each session (as seen in Fig. 4b), due both to the larger differences between the reactivation patterns and the stimulus-evoked response patterns early in the session, and to the increased number of reactivations per trial (and consequent model iterations) early in the session (Fig. 2b). Indeed, this model accurately captured all future stimulus response patterns and the more rapid change in patterns in early trials (Fig. 4f and 4g). Further, for any given session, the projection of these stimulus-evoked response patterns exhibited small fluctuations on the order of several trials (Fig. 4f). By high-pass filtering the actual and modeled stimulus-evoked responses, we found that our model was even able to capture these trial-to-trial fluctuations in stimulus-evoked response patterns (Fig. 4g, insets), further suggesting rapid plasticity that depends on the instantaneous content and rate of reactivations.

Our highly simplified model also captured the gradual orthogonalization of responses to S1 and S2 (Fig. 4h). As with the actual data (Fig. 3e-g), the orthogonalization in the modeled data was driven by Decrease neurons and not by Increase neurons (Fig. 4i-j, Extended Data Fig. S4d-e; neuron classes defined using actual data, Fig. 3d). Notably, as in the actual data, modeled activity of Decrease neurons decreased differentially across repeated presentations of S1 or S2, while modeled activity of Increase neurons increased similarly across S1 and S2 presentations (Fig. 4i-j). Thus, a very simple model using only reactivations can capture not only the nature but also the rate of change in stimulus-evoked response patterns, including the selective changes in stimulus tuning in a subset of stimulus-driven neurons (i.e. in Decrease neurons). In summary, stimulus reactivations during the ITI, while by definition somewhat similar to early stimulus-evoked responses, nevertheless differ in pattern from early stimulus-evoked responses. Moreover, these stimulus reactivations appear to draw stimulus-evoked response patterns towards them over the course of a session at a rate proportional to the reactivation rate (Fig. 4k). Overall, these findings suggest that unexpected stimuli induce reactivations in lateral visual cortex that may guide plasticity and orthogonalization of distinct stimulus representations.

## Discussion

We observed transient reactivations of specific salient stimuli in lateral visual cortex during periods of quiet waking in the tens of seconds following stimulus presentation, providing a bridge between studies of offline reactivation in sensory cortex^21, 23, 27^ and studies in hippocampus and elsewhere with similar observations during spatial navigation tasks^15-24^. We found that lateral visual cortex population responses to distinct stimuli became more orthogonal across repeated presentations^4^ while maintaining homeostatic levels of overall evoked activity^45-48^. Prior to this work, it remained unclear whether offline reactivations relate to pattern separation in any brain area. Existing hypotheses posited that reactivations are copies of previous experiences^1, 11, 12^, and that they serve to stabilize the pattern of activity that occurred during these experiences^1, 11, 12, 31, 49, 50^. By contrast, the patterns of stimulus-evoked reactivations that we observed following initial stimulus presentations in each session already differed in a systematic way from preceding stimulus-evoked response patterns, in that they more strongly resembled late-session stimulus-evoked patterns. Indeed, by feeding only the set of recorded reactivations that followed each stimulus into a simple model, we could predict the evolution of stimulus response patterns and response orthogonalization throughout the session, including dynamic changes in plasticity rates over time. Thus, our findings suggest that stimulus reactivations may play a more instructive role than previously appreciated in shaping and orthogonalizing neural representations of recently experienced stimuli. Causal interrogation of this hypothesis should soon be possible using emerging electrophysiological technologies that allow for simultaneous recordings of thousands of neurons^51^, thereby matching the sensitivity of our approach, while also allowing sufficiently fast closed-loop silencing of content-specific reactivations^52-54^.

Single trial optogenetic silencing of lateral visual cortex during a stimulus presentation prevented the selective increase in reactivations of that stimulus during the following tens of seconds. This demonstrates that the participation of lateral visual cortex neurons in stimulus reactivations requires prior activation of these same neurons during the stimulus. Further, these results suggest that some of the changes underlying response orthogonalization may involve local synaptic plasticity, in addition to other potential mechanisms^4^. Indeed, the model we used to accurately predict which neurons would increase or decrease their stimulus responses based on over- or under-participation in reactivations could be implemented biologically using a simple Hebbian learning rule. Specifically, neurons that over- or under-participate in stimulus reactivations may strengthen or weaken their connectivity, respectively, to other neurons in the co-activated stimulus-evoked ensemble^15, 55-57^. If so, future stimuli that activate parts of this ensemble would differentially recruit over-participating neurons (i.e. pattern completion)^57^.

Why does lateral visual association cortex orthogonalize stimulus representations? Separating the representations of distinct stimuli might prevent overgeneralization during the process of associating objects with outcomes. Offline reactivations might accelerate orthogonalization in the absence of frequent experience with each stimulus (which is unlikely for most salient stimuli in nature), while also helping to link visual stimuli with other reliably co-occurring, multimodal stimuli during a given experience. It remains unclear how early stimulus reactivation patterns could resemble late stimulus-evoked patterns. This may reflect pre-existing biases in local cortical connectivity that result in a manifold of preferred cortical activity patterns^18^. Feedforward input during presentation of our visual stimuli may drive activity patterns that initially stray somewhat from this manifold, with experience-dependent plasticity then pulling the sensory-evoked patterns back to the nearest location on this manifold^58^.

In summary, our study indicates that local stimulus-evoked activity patterns in lateral visual cortex give rise to reverberating reactivations of similar but not identical activity patterns in the tens of seconds following the stimulus. Stimulus-evoked patterns gravitate towards reactivated patterns at a pace proportional to the reactivation rate and to the residual difference between evoked and reactivated patterns. This brings forth the idea that, more generally, offline reactivations may actively *re-organize* sensory evoked response patterns to enhance separability of population responses during distinct experiences^59, 60^ while also potentially supporting pattern completion^61^, stabilization^49^ and associative learning^23^.

We used multinomial logistic regression (see Extended Data Fig. S1a and Methods) to decode the content of moments of synchronous spontaneous activity across stimulus-driven neurons during the inter-trial interval. This resulted in estimates of whether the spontaneous pattern resembled the S1-evoked pattern (S1 reactivation probability, top, green), the S2-evoked pattern (S2 reactivation probability, top, red), or patterns observed during non-synchronous moments within the inter-trial interval. **e**, Mean pupil area (top: normalized to maximum across the session; middle: relative change) and brain motion plotted surrounding the onset of classified reactivations (i.e. those with reactivation probability > .75) across 7 mice (two-tailed paired t-test). **f**, Left: example stimulus-evoked response from a single session for all stimulus-driven neurons across lateral visual cortex. Right: comparison of mean stimulus-evoked activity between LI, POR, P, and LM (n = 4, one-way ANOVA, Tukey HSD corrected). **g**, Same as **f** but for stimulus reactivation activity (n = 4, one-way ANOVA, Tukey HSD corrected). In all figures, the mean ± SEM is shown as error bars. n.s.: not significant, p > .05; * p < .05; ** p < .01, *** p < .001, **** p < .0001.

---

**Extended Data Figure 1.**
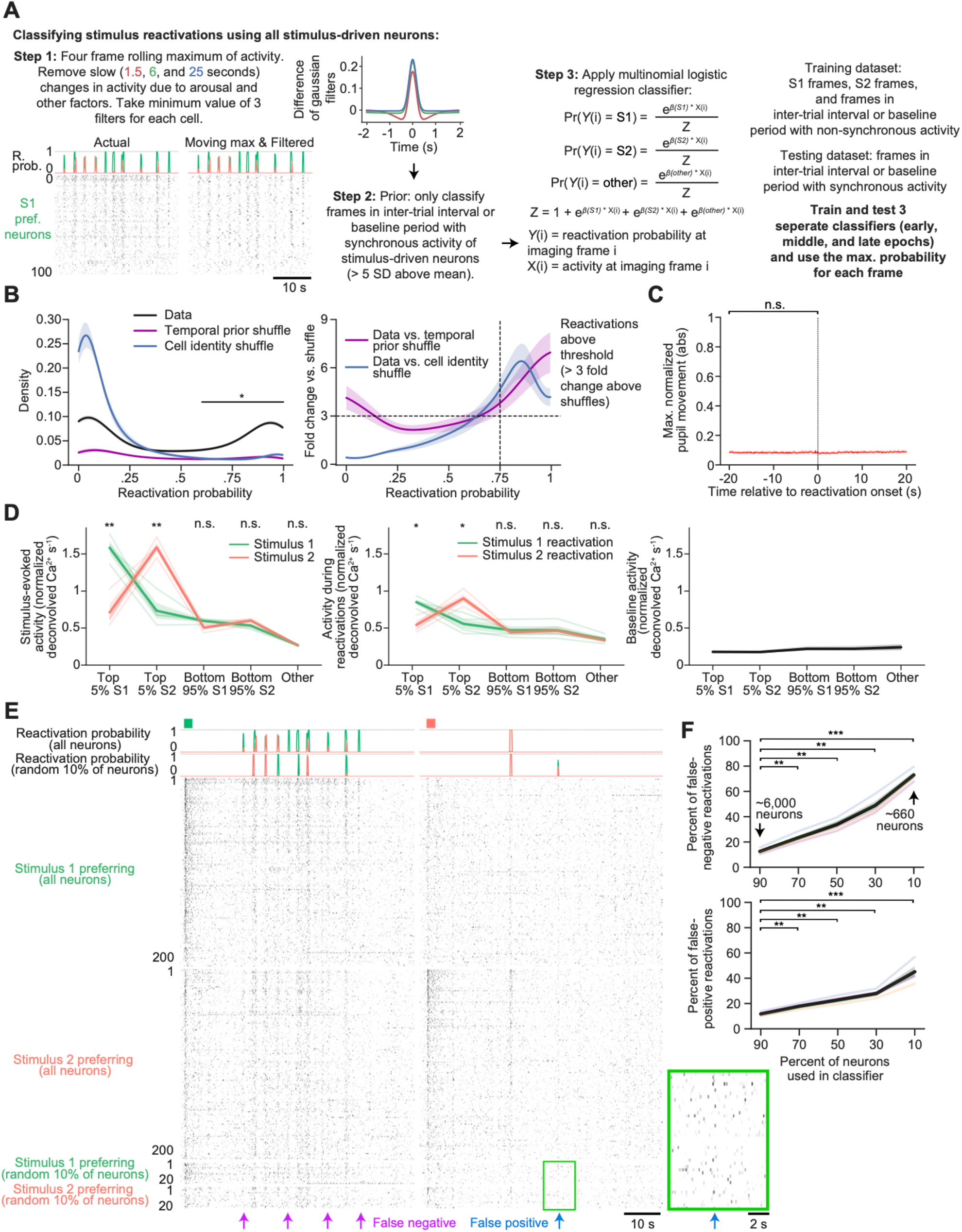
Classifying and characterizing stimulus reactivations. **a**, Brief summary of method for classifying reactivations (for additional details, see main text and Methods). Left (Step 1): we assume that the classifier should identify transient synchronous reactivations lasting at least several hundreds of milliseconds^1, 12, 16, 21, 23^ and estimate population activity patterns using the rolling maximum activity of each cell across ∼380 ms. We then remove slow changes in ongoing Ca^2+^ activity by using 3 difference-of-Gaussian filters to high-pass filter activity changes at time scales of 1.5, 6, and 25 seconds. Middle (Step 2): we define S1 or S2 stimulus reactivations during the inter-trial interval (in which the mouse passively views a mean-luminance blank screen) as epochs of synchronous activity lasting hundreds of milliseconds across neurons driven by stimulus S1 or S2, respectively. To focus on synchronous events, we use a binary prior such that we only classify reactivation pattern content during epochs in which the ongoing activity trace averaged across the top stimulus-driven neurons exceeds 5 standard deviations above the mean. Right (Step 3): we then apply multinomial logistic regression to epochs specified by the temporal prior. We train the classifier on time points that occur during all S1 trials, all S2 trials, and all time points during inter-trial intervals and during the baseline period that do not exhibit synchronous activity of stimulus-driven neurons (temporal prior = 0). We then apply the classifier to all time points with synchronous activity of stimulus-driven neurons during inter-trial intervals and during the baseline period prior to any stimulus presentations (temporal prior = 1). This results in matching probability estimates that the pattern at each time point matches the S1-evoked response pattern or S2-evoked response pattern (i.e. S1 or S2 reactivation probabilities), or ‘other’ patterns, with the sum of these probabilities equaling 1 for each time point. **b**, Left: distribution of reactivation probabilities of the classifier trained using the actual data and trained using data after shuffling using one of two different methods. The first shuffle method defines the temporal prior using an equal number of randomly selected neurons instead of only stimulus-driven neurons, and the second shuffle method randomly assigns the stimulus-evoked response of S1 or S2 neurons (n = 7 mice including 4 mice in which we ran the standard experimental protocol, as well as 3 additional mice from optogenetic silencing experiments in Figure 2d-g; one-way ANOVA, Holm-Bonferroni corrected, all data points below the significance line indicate classifier probabilities that differ significantly from both shuffled versions). Right: fold change in density of each reactivation probability using the actual data as compared to each shuffle (n = 7 mice). We defined reactivation events as those with a peak probability greater than 0.75, as they were greater than three times more common as compared to shuffled data. **c**, Peak-normalized pupil movement (absolute change in movement) plotted surrounding the onset of classified reactivation events (n = 7 mice, two-tailed paired t-test). **d**, Left: mean S1-evoked activity (green) or S2-evoked activity (red) for the top 5% and bottom 95% of S1- or S2-driven neurons and for other neurons lacking stimulus-evoked Ca^2+^ activity (n = 4 mice, two-tailed paired t-test, Holm-Bonferroni corrected). Thin lines: individual mice. Thick lines: mean across all mice. Middle: same as left but for mean Ca^2+^ activity during reactivation events (n = 4, two-tailed paired t-test, Holm-Bonferroni corrected). Right: baseline Ca^2+^ activity (in the 0.5 hours prior to any stimulus presentations) for the top 5% and bottom 95% of S1- or S2-driven neurons and for other neurons lacking stimulus-evoked Ca^2+^ activity. **e**, Raster plot of ongoing deconvolved Ca^2+^ activity of the top stimulus-driven neurons during and following an example S1 stimulus presentation (green square) and an example S2 stimulus presentation (red square), using all neurons or using a random 10% of neurons (see lower raster). Classification of stimulus reactivations using a random 10% of neurons results in several false positive (blue arrows) and false negative (magenta arrow) classification errors when compared to using all neurons. Inset at right: expanded view of data from green rectangle, illustrating a false-positive classification using a random 10% of neurons. **f**, Percent false negative (top) or false positive (bottom) classifications of reactivations relative to reactivations classified using all neurons, plotted as a function of the percent of randomly selected neurons used in the classifier (n = 4, two-tailed paired t-test, Holm-Bonferroni corrected).

**Extended Data Figure 2.**
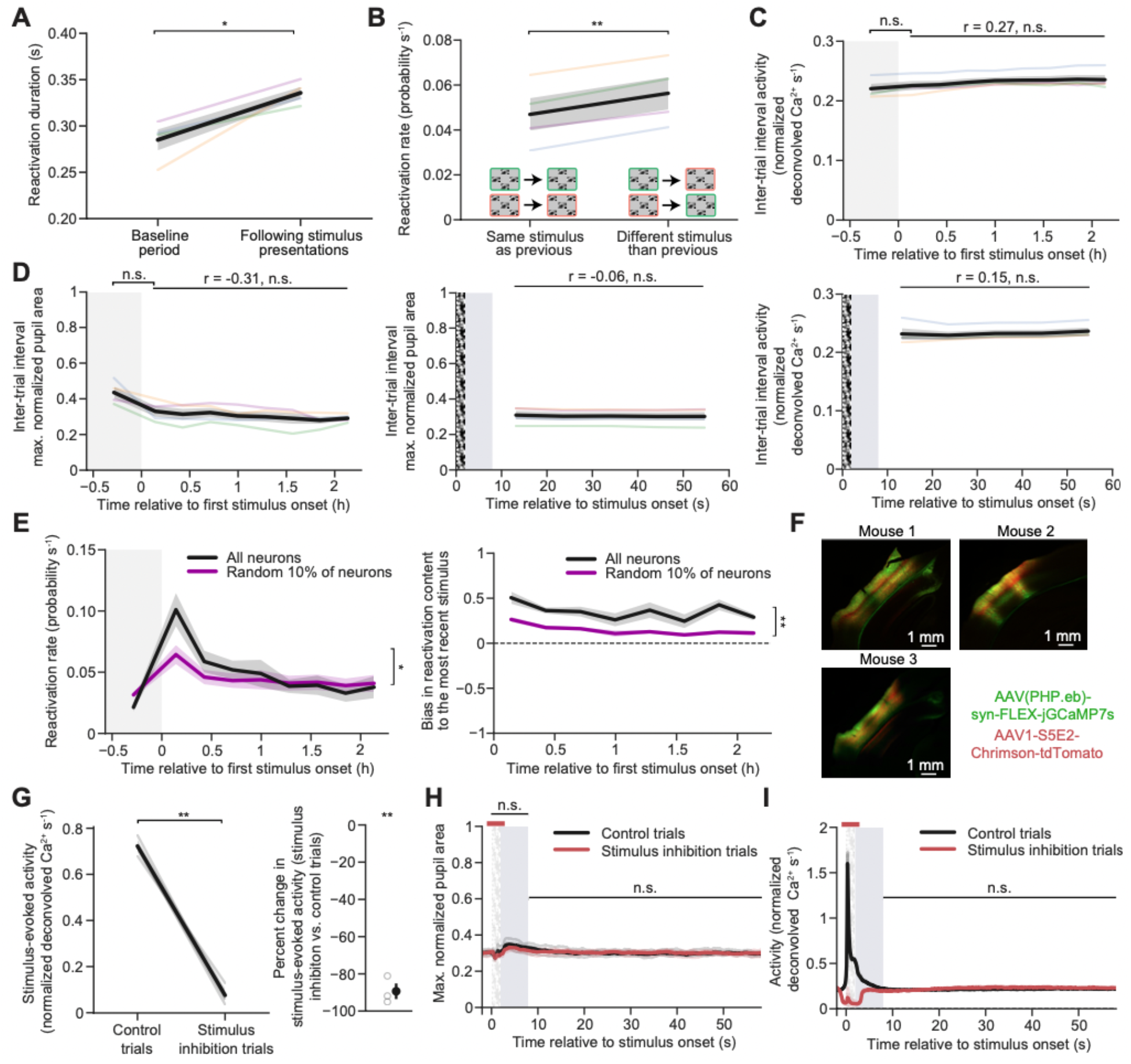
Stimulus-evoked Ca^2+^ activity and arousal state are constant during the inter-trial interval and throughout the session. **a**, Reactivation duration during the baseline period before any stimulus presentations vs. during the inter-trial intervals between stimulus presentations (n = 4, two-tailed paired t-test). **b**, Stimulus reactivation rates when the stimulus on the preceding trial was different vs. when it was the same as on the current trial (n = 4, two-tailed paired t-test). **c**, Left: Ca^2+^ activity during the baseline period before any stimulus presentation (shaded region) and during inter-trial intervals between stimulus presentations (n = 4, two-tailed paired t-test, two-sided linear least-squares regression). Right: Ca^2+^ activity throughout the inter-trial interval (n = 4, two-sided linear least-squares regression). Shaded region: excluded portion of inter-trial interval. White-noise bar: stimulus presentation. Colored lines: individual mice. Black line: mean across mice. **d**, Left: peak-normalized pupil area during the baseline period before any stimulus presentation and during inter-trial intervals between stimulus presentations (n = 4, two-tailed paired t-test, two-sided linear least-squares regression). Right: peak-normalized pupil area during the inter-trial interval (n = 4, two-sided linear least-squares regression). Shaded region: excluded portion of inter-trial interval. White-noise bar: stimulus presentation. Colored lines: individual mice. Black line: mean across mice. **e**, Left: reactivation rates during the baseline period before any stimulus presentation (shaded area) and during the inter-trial intervals between stimulus presentations, using all neurons vs. a random 10% of neurons (n = 4, two-tailed paired t-test between mean of traces excluding the baseline period). Right: reactivation content bias during stimulus presentations across the session using all neurons vs. a random 10% of neurons (n = 4, two-tailed paired t-test between mean of traces). **f**, Coronal sections of visual cortex displaying virally-mediated expression of Cre-dependent jGCaMP7s in glutamatergic neurons (green, in Emx1-Cre mice) and Chrimson in parvalbumin interneurons (red, S5E2 enhancer) in 3 mice. Local injections ensured targeted expression of Chrimson throughout lateral visual cortical areas in all 3 mice. **g**, Left: stimulus-evoked Ca^2+^ activity during control vs. stimulus inhibition trials (n = 3, two-tailed paired t-test). Right: percent reduction in Ca^2+^ activity on stimulus inhibition trials compared to control trials (one-sample t-test vs. 0). For stimulus inhibition trials, we pulsed 10 mW of red light for 4 ms at 16 Hz from 1 second before stimulus onset to 1 second after stimulus offset. **h**, Peak-normalized pupil area during stimulus presentation and during the inter-trial interval for control vs. stimulus inhibition trials (n = 3, two-tailed paired t-test between mean of traces during the specified inter-trial interval or stimulus period plus the period immediately following the stimulus, Holm-Bonferroni corrected). Red horizontal bar at top indicates timing of optogenetic silencing. Noise bars indicate time of visual stimulus. Gray shaded area indicates post-stimulus period excluded from reactivation analyses. **i**, Ca^2+^ activity during stimulus presentation and during the inter-trial interval for control vs. stimulus inhibition trials (n = 3, two-tailed paired t-test between mean of traces during the specified inter-trial interval). Red horizontal bar at top indicates timing of optogenetic silencing. Noise bars indicate time of visual stimulus. Gray shaded area indicates post-stimulus period excluded from reactivation analyses.

**Extended Data Figure 3.**
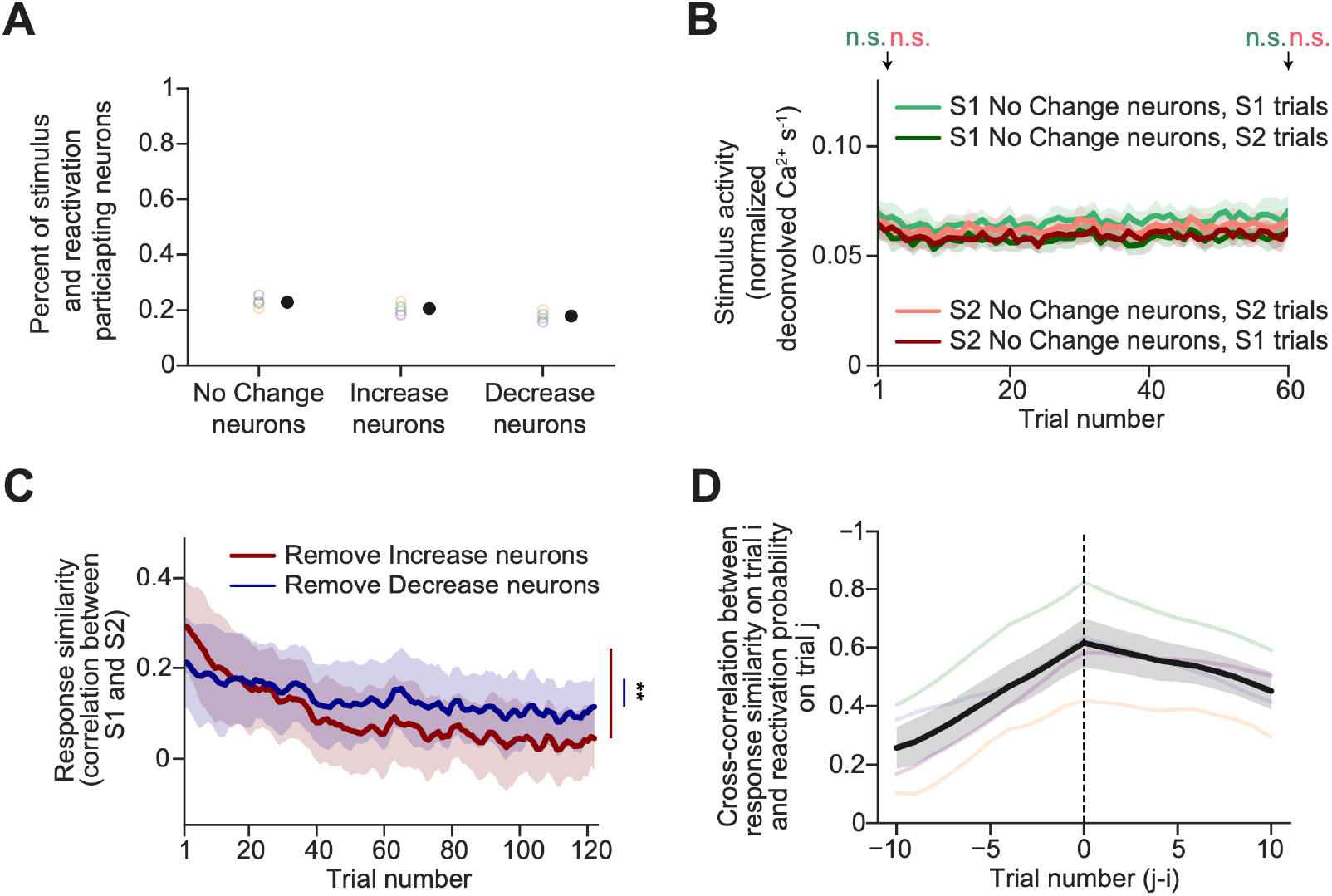
Characterization of No Change, Increase, and Decrease neurons. **a**, Percent of neurons that are grouped as No Change, Increase, or Decrease neurons. **b**, The evoked Ca^2+^ activity of No Change neurons, split by neurons whose response does not change to S1 (light green) or S2 (pink). Also shown are the responses of these two groups of neurons to the other stimulus (S2 [dark green] and S1 [dark red], respectively; n = 4, two-tailed paired t-test, Holm-Bonferroni corrected). **c**, Response similarity (running correlation between response patterns during neighboring S1 and S2 trials), plotted in the same manner as the mean trace in **Fig. 3c**, but calculated after removing all Increase neurons (red) or Decrease neurons (blue) (n = 4, two-tailed paired t-test). Traces were smoothed with a 3-trial sliding average. **d**, Cross correlation between response similarity and reactivation probability (n = 4).

**Extended Data Figure 4.**
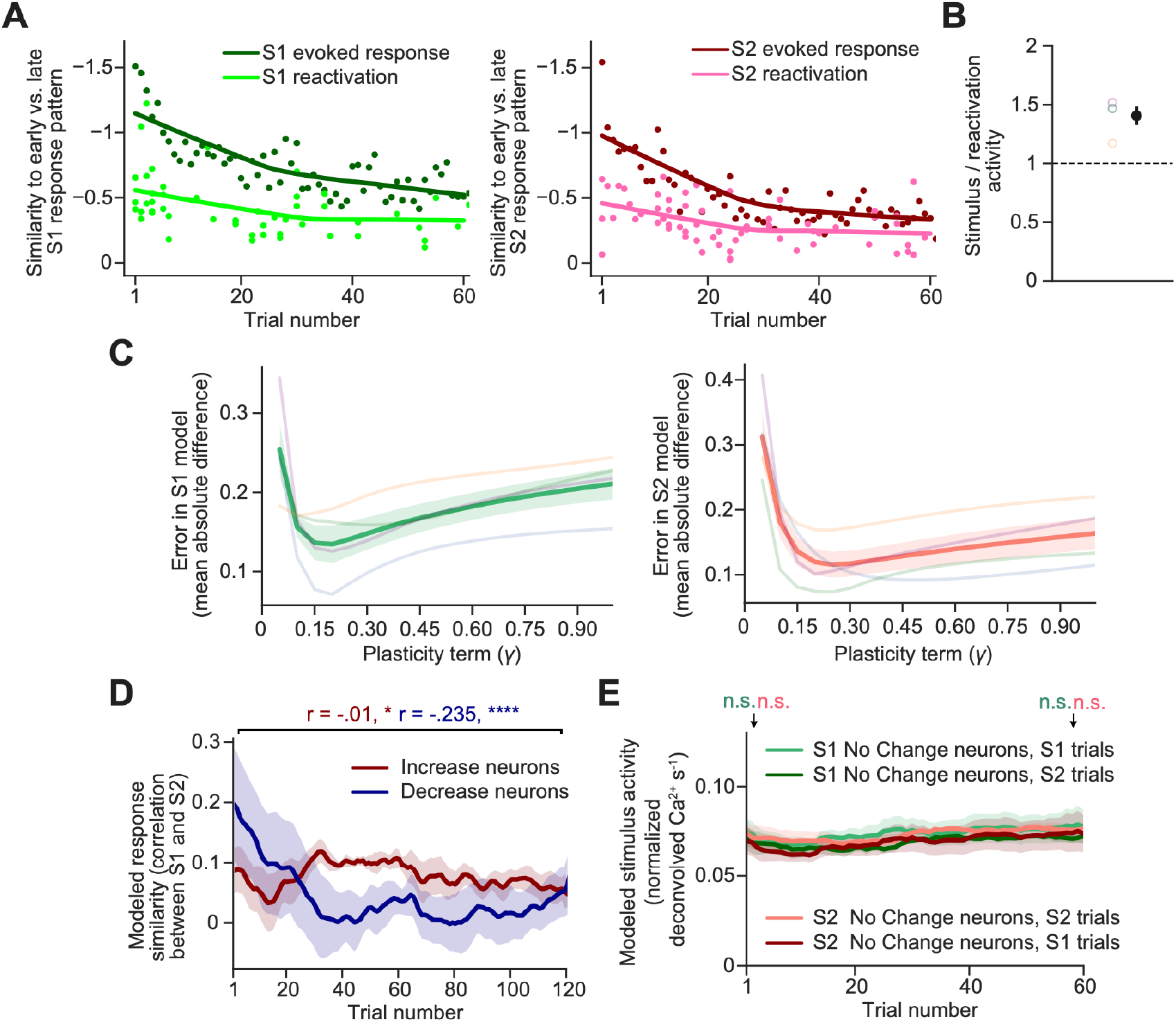
Modeling future stimulus responses using reactivations. **a**, For each trial, we projected single-trial response patterns (during S1 or S2) and stimulus reactivations during the inter-trial interval (S1R or S2R) onto the axis between early and late stimulus-evoked response patterns within a session (see **Fig. 4a-b** for additional graphical details). Here, data from a typical example session is shown. **b**. Stimulus-evoked activity vs. reactivation activity, averaged across all neurons that were both stimulus-driven and that participated in reactivation events early in the session (see Methods). See also **Extended Data Fig. 1d. c**, We parametrically varied the plasticity variable γ and measured the error in the modeled stimulus-evoked response patterns vs. actual stimulus-evoked response patterns (mean of the absolute difference between actual and modeled data). **d**. As in **Fig. 3e**, response similarity (running correlation between responses of neighboring S1 and S2 trials) for modeled stimulus response patterns, calculated separately using only Increase or Decrease neurons (as defined using actual data; n = 4, two-tailed least squares linear regression, Holm-Bonferroni corrected). **e**, Mean stimulus-evoked activity using modeled data, averaged across S1 No Change neurons or S2 No Change neurons (as defined using actual data; cf. **Fig. 3d and Extended Data Fig. S3b**) (n = 4, two-tailed paired t-test, Holm-Bonferroni corrected).

## Methods

### Data reporting

No statistical methods were used to predetermine sample size. Experiments did not involve experimenter blinding and were not randomized.

### Mice

All animal care and experimental procedures were approved by the Beth Israel Deaconess Medical Center Institutional Animal Care and Use Committee. Animals were group housed before surgery and singly housed post-surgery. Animals were provided ad libitum access to standard mouse chow and water for all experiments. Animals were on a 12-hour dark/light cycle. We used 4 mice (3 female, 1 male) for standard experiments and 3 mice (2 female, 1 male) for experiments that also involved optogenetic inhibition. All animals were adult (> P56) transgenic (Emx1-cre) mice. All experiments were performed during the light cycle.

### Behavioral training

Mice were first habituated to head fixation for 4-7 days. On the first day, mice were head-fixed for 1 hour and allowed to run on the wheel in the dark. On following days and during imaging experiments, mice were head-fixed and the wheel was also fixed such that mice could not run, but could adjust their posture laterally. We also presented a gray screen using a Dell 60 Hz LCD monitor positioned on the right side of the mouse for habituation purposes. Mice were progressively habituated for an additional 30 minutes each day. We continued these daily habituation sessions until the point where the mice remained calm, and their eyes remained clear without physical indications of stress for two hours.

We then began daily recording sessions. Each day, mice were first were presented with a gray screen for ∼0.5 hours. Following this baseline period, mice were presented with one of two checkerboard patterns (Stimulus 1 [S1] or Stimulus 2 [S2]), with each square in the checkerboard containing the same binarized white noise movie^62^. Each stimulus lasted for 2 seconds followed by a 58 second inter-trial interval during which a gray screen was shown. Binarized white noise visual stimuli and the mean luminance gray screen during the inter-trial interval were luminance matched. Each session lasted ∼3 hours and consisted of the 0.5-hour baseline period followed by 64 presentations of each of the two stimuli in a random order. The visual stimulation spanned a large part of the monocular region of the mouse visual field, from ∼-3° to 93° in azimuth and ∼- 42° to 42 in elevation.

### Surgical procedures

We followed previous surgical procedures for cranial windows^63^. Briefly, in anesthetized mice, we performed a 3-millimeter circular craniotomy with a dental drill on the left lateral visual cortex (centered at AP: -4.6 mm, ML: -4.35 mm with respect to Bregma). We placed a 3-mm circular clear window (glued to a 5-mm clear window on top with edges that rest on the thinned skull) onto the brain surface. We fixed the window in place with C & B Metabond (Parkell). Prior to performing viral injections of AAV(PHP.eb)-syn-FLEX-jGCaMP7s^34^ (Addgene), we first waited approximately 1 week for mice to recover, brain edema to decline, and blood to clear from below the cranial window. We then removed the window under anesthesia and performed 18 injections (33-100 nl each; depths: 200, 350, and 500 μm; speed of injection: 10-30 nl/min), evenly spaced throughout the exposed 3-mm diameter circular brain surface, at various dilutions for each mouse (e.g. 1:0, 1:5, 1:5, and 1:10 for the 4 mice in Figure 1; jGCaMP7s:saline). We then replaced the window with a new one and waited for mice to recover for at least 1 week. For optogenetic inhibition studies, the same procedure was performed but we instead injected a mixture of jGCaMP7s, S5E2-Chrimson, and saline (at a ratio of 0.75 : 3 : 3.75).

### Retinotopy

We used brief epifluorescence imaging of jGCaMP7s signals to obtain retinotopic maps of local preference for specific locations in visual space in order to identify several visual areas: V1: primary visual cortex; LM: lateromedial; POR: postrhinal; LI: laterointermediate; P: posterior. We presented low contrast vertical gratings displayed at 1 of 4 different retinotopic locations. A 470 nm LED passed through a long-pass emission filter (500 nm cutoff). Images were collected using a EMCCD camera. To determine which neuron(s) belong to which visual region (P, LI, POR, LM), we manually drew region boundaries (after alignment and resizing to a reference atlas^64, 65^) using the roipoly function in MATLAB. For each neuron, we determined its center of mass for each spatial mask and assigned its location to P, LI, POR, or LM accordingly.

### Two-photon calcium imaging

We measured Ca^2+^ activity using two-photon microscopy. We used an Insight X3 laser from Spectra-Physics to excite at 920 nm (20-40 mW). Imaging was performed using an Olympus 10x water immersion objective (0.6 NA; 796 × 512 pixels spanning an area of ∼2000 × 1500 μm) on a resonance-scanning two-photon microscope (Neurolabware and Optotune; near-simultaneous imaging of three planes each spaced >20 μm apart, 31.25 Hz total imaging rate for a sampling rate of 10.42 Hz for neurons at each plane). We imaged layers II-IV (approximate depth of 180-380 μm from the brain surface) of lateral visual cortical regions (LI, POR, P, LM, and sometimes very lateral V1). We collected data in 34-minute runs (1 baseline run and 4 stimulus presentations runs) in a dark and quiet room with limited entry. Mice were imaged on consecutive days or every other day over several weeks (maximum duration less than one month).

### Image processing and source extraction

We used suite2p to align, register, detect cell masks, extract Ca^2+^ fluorescence traces, and deconvolve these traces^37^. Briefly, we used non-rigid motion correction in blocks of 128 × 128 pixels and registered each chunk for each frame to a reference. We defined ‘brain motion’ (Fig. 1e) as the absolute amount of shift when aligning each image. For cell detection, suite2p decomposes the data into a low-dimensional form and clusters to find regions of interests (ROIs) consisting of correlated pixels. Mean fluorescence intensities are then extracted from each ROI and the surrounding neuropil (excluding other ROI masks). For deconvolution, we first corrected for neuropil contamination by subtracting the mean surrounding neuropil fluorescence from each ROI fluorescence trace with a neuropil coefficient (scale factor) of 0.7. Fluorescence traces were then corrected for long time scale drift by subtracting a 60-second sliding window median filter. The OASIS algorithm was then applied to this corrected fluorescence trace to obtain non-negative spike deconvolution^66^. For all analyses, we used peak-normalized, deconvolved Ca^2+^ activity. Specifically, we normalized each cell’s deconvolved activity trace separately, dividing all values by the mean of the top 1% of values. To avoid considering duplicate masks belonging to the same neuron imaged in different planes, we first aligned the planes relative to each other using displacement field estimates (imregdemons in MATLAB). After alignment, we computed the correlation of the extracted fluorescence traces across the entire recording on pairs of neuron masks that exhibited any overlap. If a pair of masks exhibited greater than 25% correlation (empirically determined) in their fluorescence traces, one of the masks was removed from further analyses.

### Pupil tracking

An IR camera was positioned below the monitor to record the right eye of each mouse. To extract the pupil size, we manually created a mask around the eye and selected the center of the pupil. This was then fit using the starburst pupil detection algorithm from the openEyes toolkit and a ransac algorithm. Rare frames with low-quality images of the eye were replaced with interpolated data.

### Stimulus-driven neurons and neurons that participated in reactivations

Neurons with significant evoked activity during S1 or S2 were determined using a non-parametric test (Wilcoxon rank-sum test). Normalized deconvolved Ca^2+^ activity for each neuron in the 2 s before stimulus presentation was compared to activity during the 2-s stimulus presentation periods. Neurons with significantly (p < .01) increased activity for S1 or S2 were defined as stimulus-driven neurons. We used a similar test to determine neurons with significantly increased activity during S1 or S2 reactivations. Normalized deconvolved Ca^2+^ activity for each neuron, averaged across the duration of the S1 or S2 reactivations, was compared to activity during a similar duration (separately for each reactivation duration) prior to each reactivation (excluding previous time points that included reactivations). Neurons with significantly (p < .01) increased activity were defined to participate in at least some stimulus reactivations.

### Classifying stimulus reactivations

For classification of transient stimulus reactivation patterns, we used the normalized deconvolved Ca^2+^ activity of S1 and S2 stimulus-driven neurons. We assumed that the classifier should identify transient synchronous reactivation events lasting at least several hundreds of milliseconds (4 frames or ∼380 ms), a time scale roughly similar to that in previous studies^1, 12, 16, 21, 23^. Thus, for epochs used to assess the presence of reactivations, we estimated population activity patterns using the rolling maximum activity of each cell across ∼380 ms. We then removed slow changes in activity. To remove slow changes in activity on the order of ∼1.5 s, ∼6 s, and ∼25 s (4^2^/ sampling rate, 4^3^/ sampling rate, and 4^4^/ sampling rate, respectively; sampling rate for each activity trace: 10.42 Hz), we built three difference-of-Gaussian filters by subtracting each of the three broad Gaussians separately from the narrow Gaussian. We high-pass filtered the normalized deconvolved Ca^2+^ activity of S1 and S2 stimulus-driven neurons using each of the three difference-of-Gaussian filters. For each time point, we took the minimum value across the three filtered traces to maximally remove slow changes in activity. This resulted in a filtered normalized deconvolved Ca^2+^ activity trace for S1 and S2 stimulus-driven neurons that removes slow changes in activity and preferentially preserves rapid, transient synchronous activity.

We then built a prior for synchronous activity of stimulus-driven neurons (i.e. those driven by S1 and/or S2) at each time point. Because the top 5% of stimulus-driven S1 or S2 neurons contained the most reliable information about stimulus identity (see Extended Data Fig. 1d), we found time points during the inter-trial interval and baseline period where the average activity across the top 5% of stimulus-driven neurons was greater than 5 standard deviations above the mean. These time points were used as a prior for synchronous activity of stimulus-driven neurons, and we only classified the content of stimulus reactivations within these time points (see also Extended Data Fig. 1a).

To classify the identity of synchronous activity of stimulus-driven neurons during the inter-trial interval and baseline period, we used multinomial logistic regression, an extension of logistic regression. We excluded the 6 seconds immediately following stimulus offset during the inter-trial interval from all classifier training and testing, to allow Ca^2+^ activity to return close to baseline (Extended Data Fig. S2i). We trained the classifier with three different classes of time points: all time points during S1 presentation, all time points during S2 presentation, and all time points during the inter-trial interval and baseline period but excluding time points with synchronous activity of S1 or S2 neurons mentioned above and 6 seconds post stimulus offset (‘other’). We then tested on all time points during the inter-trial interval and baseline period with synchronous activity of S1 or S2 neurons excluding the 6 seconds post stimulus offset. This resulted in matching probability estimates that the pattern at each time point matches the S1-evoked response pattern or the S2-evoked response pattern (i.e. S1 or S2 reactivation probabilities), or ‘other’ patterns, with the sum of these three probabilities equaling 1 for each time point. Individual events with high synchronous activity could in some cases contain both S1 and S2 reactivations.

S1-evoked and S2-evoked patterns changed across time with repeated presentations (see Fig. 3 and 4b). To account for changes in stimulus representations across a session, we built three separate classifiers so as not to bias our final classification towards detecting reactivation patterns that were similar only to early or late periods within a session. We split the trials into three equal chunks (from early, middle, and late periods) and trained the classifier on each chunk separately. We then applied each classifier to all eligible time points during the ITI (or during the baseline period). For each time point, we compared the performance of early, middle, and late classifiers, and selected the probabilities from the classifier with the highest total matching probability (summed across S1 and S2). If multiple classifiers had the same total matching probability for a given frame, we used the classifier trained on data including those from nearby stimulus presentations. Note that all main results held when we instead used a single classifier trained on stimulus responses during all trials (not shown).

### Quantifying reactivation rate and bias

Reactivation rates of a given stimulus were calculated as the summed probability of reactivations of that stimulus per second. We excluded from these analyses any time points where the mouse was attempting to walk/run or when pupil motion was high (>6% displacement relative to the diameter of the eye). Reactivation bias towards S1 was calculated as the difference between the rates of S1 and S2 reactivations, divided by the sum of S1 and S2 reactivation rates. Reactivation bias towards S2 was calculated as the difference between the rates of S2 and S1, divided by the sum of S1 and S2 reactivation rates. Reactivation duration was calculated as the entire time of each reactivation (i.e. the duration of contiguous time points where the matching probability exceeded 0).

### Classifying stimulus reactivations using subsets of neurons

To train and test the classifier using subsets of neurons, we randomly selected a fraction of the total imaged neurons (from 10-90%, increasing 10% each time) and re-ran the same classifier. This was performed for 10 iterations for each subset percentage. The rate of false-negative and false-positive stimulus reactivations using subsets of total imaged neurons (Extended Data Fig. S1f) and the reactivation rate and bias using a random 10% of neurons (Extended Data Fig. S2e) was averaged from the result of 10 iterations.

### Optogenetic inhibition

For optogenetic inhibition of visual stimulus-evoked activity, photostimulation of Chrimson-expressing parvalbumin interneurons began 1 second before stimulus onset and ended 1 second after stimulus offset on inhibition trials, which occurred on a random 50% of all trials. The photomultiplier tube (PMT; H11706-40; Hamamatsu) was gated at the beginning of each frame for 6 ms to protect it from the LED light that was delivered for the first 4 ms of each frame. As a result, approximately 19% of each frame during stimulation was blanked, but allowed for simultaneous imaging of jGCaMP7s in the majority of the field of view during photostimulation (16 frames/second). A 617 nm LED (10mW, Thor labs; M617L3) controlled by a driver (Thorlabs; T-Cube) was used.

### Response metric

For all analyses in Figures 3 and 4, we used all neurons that were considered stimulus-driven by either S1 and/or S2, and that participated in at least some stimulus reactivations (see definitions above). To calculate the running Pearson’s correlation between S1 and S2 stimulus-evoked patterns, we took the mean normalized deconvolved Ca^2+^ activity for all neurons during the entire stimulus or reactivation between all pairs of neighboring S1 and S2 trials. Thus, running correlations were solely computed on trials in which an S1 trial was preceded by an S2 trial or in which an S2 trial was preceded by an S1 trial. For plotting the stimulus-evoked activity and running correlation across trials (Fig. 3a, Fig. 3c-g, Extended Data Fig. S3b-c, Fig. 4h-j, Extended Data Fig. S4d-e), we smoothed the traces by taking a moving mean of 3 trials.

### Grouping neurons

To group neurons into sets based on their changes in stimulus-evoked activity from early to late in a session, we first calculated the percent difference in normalized deconvolved Ca^2+^ activity for S1 or S2 stimulus-driven and reactivation-participating neurons between the mean of the first 10 trials and the mean of the last 10 trials. We plotted this as a distribution for each day for each mouse. No Change neurons were classified as neurons whose percent change in activity was within 0.5 standard deviations of the population mean change in activity across all cells. Increase neurons were classified as neurons whose percent change in activity was greater than 1 standard deviation above the population mean. Decrease neurons were classified as neurons whose percent change in activity was less than 1 standard deviation below the population mean.

### Cross correlation analysis

For the display of traces, correlation and cross correlation analyses in Figure 3h-i and Extended Data Figure S3d, we smoothed the traces by taking a moving average of 8 trials. We also used this smoothed moving average of 8 trials to calculate correlations and cross correlations related to estimates of response similarity.

### High-pass filtering

To high-pass filter time series (i.e. to remove slow trends from the 8 trial moving average traces for analyses in Fig. 3i, right and Fig. 4g, insets), we used a 2^nd^ order Butterworth high-pass filter with a normalized critical frequency of .05 where 1 is the Nyquist frequency. This was sufficient to allow for short-timescale fluctuations to pass while removing the roughly exponential decay in the traces. Because the exponential decay was large early in the time series, the filter was unable to fully remove this, and we thus omitted the first 5 trials for related analyses.

### Vector analysis

To calculate a vector that defined stimulus responses from early to late (Figure 4a), we took the mean stimulus response of the first 3 trials of S1 or S2 (early response), and of the last 3 trials (late response), for each of the ‘N’ stimulus-driven and reactivation-participating neurons. We then calculated the N-dimensional vector that defined the evolution from early to late responses as the late response vector minus the early response vector, separately for S1 and S2. We then projected the single-trial mean responses of S1 trials onto this S1 vector (scalar projection, the dot product with the S1 vector divided by the norm of the S1 vector), and of S2 trials onto this S2 vector. We also projected single S1 reactivations onto this S1 vector, and S2 reactivations onto this S2 vector. Thus, population response patterns that were more similar to the late response pattern than the early response pattern exhibited projection values that were more positive.

### Normalizing reactivation responses

To scale the reactivation responses to have the same magnitude as the stimulus responses, we calculated the mean deconvolved activity across stimulus-driven and reactivation-participating neurons, averaged across all reactivations and separately across the first 20 trials. We then divided the mean stimulus response by the mean reactivation response to obtain the scale factor for each session, and then averaged across sessions for each mouse. We then averaged across all mice to obtain a single scale factor that we applied to all sessions and mice.

### Modeling stimulus responses using reactivations

To model future stimulus responses using reactivations, we used the actual mean response of stimulus-driven and reactivation-participating neurons during the first S1 or S2 trial. For each subsequent inter-trial interval, we iteratively estimated a modeled S1 or S2 response as the sum of the previous trial’s estimated response and the difference between the scaled S1 or S2 reactivation pattern that occurred during the inter-trial interval and the current S1 or S2 response pattern, multiplied by a single plasticity value. We parametrically varied the plasticity value such that the error was lowest and chose a single value that we applied to all sessions and mice. We calculated the error as the mean of the absolute difference between modeled and actual S1 or S2 projections (see e.g. Fig. 4f), averaged across all sessions and mice. This update to the estimate was applied for each reactivation event in each inter-trial interval. Thus, a greater number of reactivations during a given inter-trial interval will lead to more iterations of the model occurred to update the predicted S1 or S2 response, and thus to a faster instantaneous trial-to-trial learning rate.

### Data analysis

All analysis was performed using custom scripts in MATLAB and Python. In all figures, the mean ± SEM is shown. n.s.: not significant, p > .05; * p < .05; ** p < .01, *** p < .001, **** p < .0001. All tests (one-tailed t-test, two-tailed t-test, Wilcoxon rank-sum or ANOVA) with multiple hypotheses were corrected for with multiple comparisons correction (Tukey HSD or Holm-Bonferroni). For statistical tests in Fig. 2f-g and Extended Data Fig. 2e,h-i, we compared the means of the two traces during the periods indicated in the figures.

## Data availability

Data is available upon request. Code is available at https://github.com/nguyenr95/reactivation.

## Acknowledgements

This project was supported by a National Defense Science and Engineering Fellowship and a Howard Hughes Medical Institute Gilliam Fellowship (NDN), NIH F32 DK112589, a Davis Family Foundation award, and a BNORC pilot grant (AL), and an NIH DP2 DK105570, DP1 AT010971-02S1, R01 MH12343, a McKnight Scholar Award, a Harvard Mind Brain Behavior Interfaculty Initiative Faculty Research Award, the Harvard Brain Science Initiative Bipolar Disorder Seed Grant, and by Kent and Liz Dauten (MLA).

## Author contributions

N.D.N. and M.L.A. conceived the project and wrote the manuscript. N.D.N. performed all experiments and analyzed all the data. A.L. designed and helped with optogenetic silencing studies. J.F. helped with surgical procedures. S.V., J.M., and J.D. produced the AAV-S5E2 Chrimson virus.

## Competing interest declaration

The authors declare no conflicts of interest.

## Notes

### Competing Interest Statement

The authors have declared no competing interest.

